# Reduced ZMPSTE24 expression leads to prelamin accumulation and development of steatosis in MASLD patients

**DOI:** 10.1101/2024.05.16.594546

**Authors:** Joseph D. Schinderle, Anqi Wu, Irina M. Bochkis

**Affiliations:** Department of Pathology, University of Pittsburgh School of Medicine, Pittsburgh, PA, 15261; Pittsburgh Liver Research Center, University of Pittsburgh School of Medicine, Pittsburgh, PA, 15261

**Author notes:** **Contact Information:** Irina M. Bochkis, Ph. D., Pittsburgh Liver Research Center, Department of Pathology, University of Pittsburgh School of Medicine, BST South, Room S446, 200 Lothrop Street, Pittsburgh, PA 15261.

## Abstract

Mutations of nuclear lamina-associated proteins LMNA and ZMPSTE24 have been associated with fatty liver. We report that the changes at the nuclear envelope we described in MASLD patients are caused by downregulation of ZMPSTE24, an enzyme that processes prelamin to mature lamin A. In addition, *Zmpste24* mutant mice develop hepatic steatosis and exhibit upregulation of p53 target genes. p53 activity is also induced in genes differentially expressed in MASLD patients. Furthermore, p53 regulates genes bound by FOXA2 in these individuals, corresponding to observations in *Zmpste24* mutants. In contrast, expression of glucose and insulin regulated genes is reduced in MASLD patients, suggesting altered glucose metabolism and insulin resistance, hallmarks of type 2 diabetes (T2D). Hence, our genomics data show that MASLD patients with severe steatosis but yet without MASH are already suffering from severe metabolic consequences and underscore the need for treatment at this stage of the disease.

## Introduction

Today, the prevalence of metabolic syndrome is on the rise and now affects 1 in three adults^1^. Metabolic syndrome is used to determine metabolic health in adults, and a patient is said to have metabolic syndrome if they have 3 of the following 5 conditions: obesity, high blood glucose levels, low levels of high-density lipoprotein cholesterol, high triglyceride levels, and high blood pressure^2^. These conditions are all associated with an increased risk of the development of Type 2 Diabetes (T2D), which affects approximately 1 in 10 people, and nearly three quarters of the US population over the age of 65^3,4^. Non-alcoholic Fatty Liver Disease (NAFLD), or Metabolic Dysfunction-associated Steatotic Liver Disease (MASLD), is extremely common for T2D patients, and is estimated that about 75% of T2D patients also suffer from MASLD^4^.

Genetic variants in nuclear lamina-related genes, including ZMPSTE24, have been associated with MASLD and changes in nuclear morphology have been observed in MASLD patients ^5 6,7^. In addition, a heterozygous mutation in ZMPSTE24 was found in a patient with severe metabolic syndrome and high hepatic triglyceride content ^8^. ZMPSTE24 is an enzyme that processes prelamin to mature lamin A. Mutations in *LMNA*, encoding the nuclear structural protein lamin A/C, due to lack of processing and accumulation of prelamin, result in nuclear lamina dysfunction and cause the premature aging syndrome Hutchinson-Gilford progeria (HGPS). Additionally, *LMNA* mutations lead to partial lipodystrophy, a condition associated with insulin-resistant diabetes, hypertriglyceridemia, renal disease, cardiomyopathy, and hepatic steatosis ^9^. Hepatocyte-specific deletion of mouse lamin A/C results in spontaneous steatosis that progresses to steatohepatitis ^10^.

We have recently shown that change in expression of lamina-associated proteins leads to dysmorphic nuclear shape and untethering of heterochromatin at the nuclear envelope, resulting in redistribution of lamina-associated domains (LADs), opening of repressed chromatin due to altered FOXA2 binding, up-regulation of genes regulating lipid synthesis and storage, culminating in fatty liver and steatosis in mouse models and MASLD patients, similar to the mechanism we reported in aging and laminopathy models ^7 11^. Laminopathy is a group of diseases caused by mutations in genes encoding nuclear lamina proteins, such as ZMPSTE24. *Zmpste24* mutant mice have irregular nuclei and develop hepatic steatosis ^12^.

Here we extend our findings, demonstrating prelamin accumulation in livers of MASLD patients due to reduced expression of ZMPSTE24 and linking these changes at the lamina to our previously described Foxa2-dependent mechanism ^7^. In addition, expression of glucose and insulin regulated genes is reduced in MASLD patients, suggesting altered glucose metabolism and insulin resistance, hallmarks of type 2 diabetes (T2D). Hence, our genomics data show that MASLD patients with severe steatosis but yet without MASH are already suffering from severe metabolic consequences and underscore the need for treatment at this stage of the disease.

## Results

### Nuclear lamina changes in female MASLD patients

We have recently shown that change in expression of lamina-associated proteins leads to dysmorphic nuclear shape and untethering of heterochromatin at the nuclear envelope, resulting in redistribution of FOXA2 binding, activation of previously repressed genes and development of hepatic steatosis in mouse models and male MASLD patients, similar to the mechanism we reported in aging and laminopathy models ^7 11^. In this study, we analyzed nuclear lamina changes in younger female MASLD patients (normal controls, mild steatosis, moderate steatosis, healthy 17-59 years old, MASLD 31-47 years old). Indeed, the nuclear shape in female MASLD patients with mild or moderate steatosis was altered (**Fig. 1a**). We also profiled lamina-associated domains (LADs) using Lamin B1 ChIP-Seq. In contrast to males, Lamin B1 ChIP-Seq signal decreases slightly in female patients with mild steatosis, and more significantly in female patients with severe steatosis (**Fig. 1b**). FOXA2 binding does not change in patients with mild steatosis and is redistributed and is repositioned in patients with moderate steatosis (**Fig. 1c**). Similar to males, lamina changes reshape FOXA2 binding in female MASLD patients.

**Figure 1.**
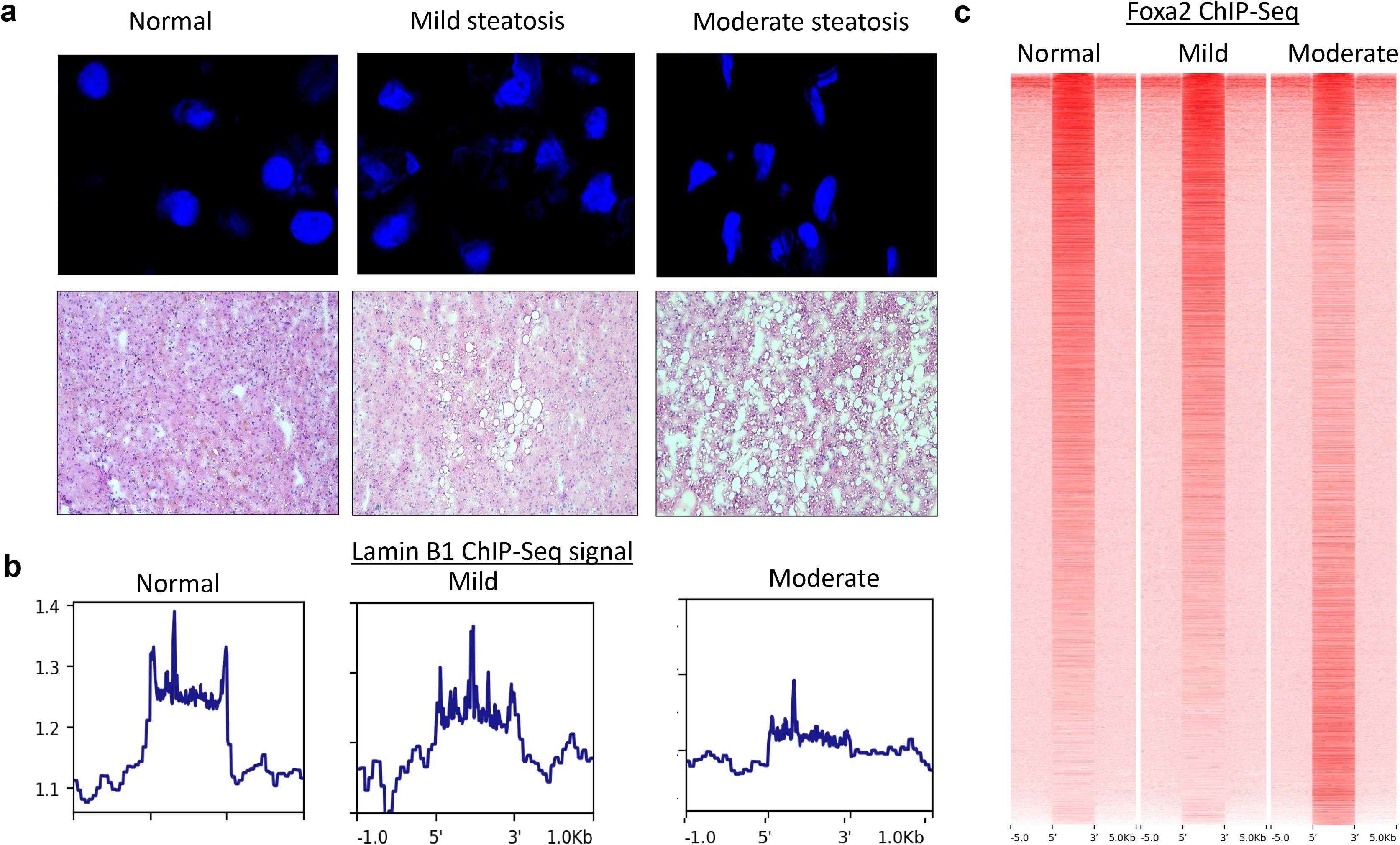
Changes at the lamina reshape Foxa2 binding in female MASLD patients during progression of the disease. **(a)** Nuclear DAPI staining (top panel) and H&E (bottom panel) staining of liver sections from normal patients (left panel) and female MASLD patients with mild (middle panel) and severe steatosis (right panel). (**b**) Lamin B1 ChIP-Seq signal (Reads Per Kilobase, per Million mapped reads, RPKM) calculated at lamina-associated domains (LADs) decreases in MASLD patients. (**c)** Heatmap of Foxa2 ChIP-Seq signal showing that Foxa2 binding in patients with mild steatosis is similar to occupancy in normal patients, but is redistributed in patients with moderate fatty liver

### p53 regulates FOXA2 targets and differentially expressed genes in MASLD patients

Next, FOXA2 binding sites in male patients with severe steatosis and female patients with moderate steatosis were mapped to genes. Ingenuity Pathway Analysis (IPA) analysis identified activation of several nuclear receptors, including RAR, CAR in both males and females and VDR and PPARα only in females. (**Fig. 2a**). Pathways associated with FOXA2 targets exclusively in livers of male MASLD patients included “Cholesterol Biosynthesis” and “Hepatic Cholestasis”, consistent with FOXA2 role we described in bile acid metabolism ^13-15^. In contrast, ‘Liver Proliferation” and “Liver Steatosis” were significant in livers of female patients.

**Figure 2.**
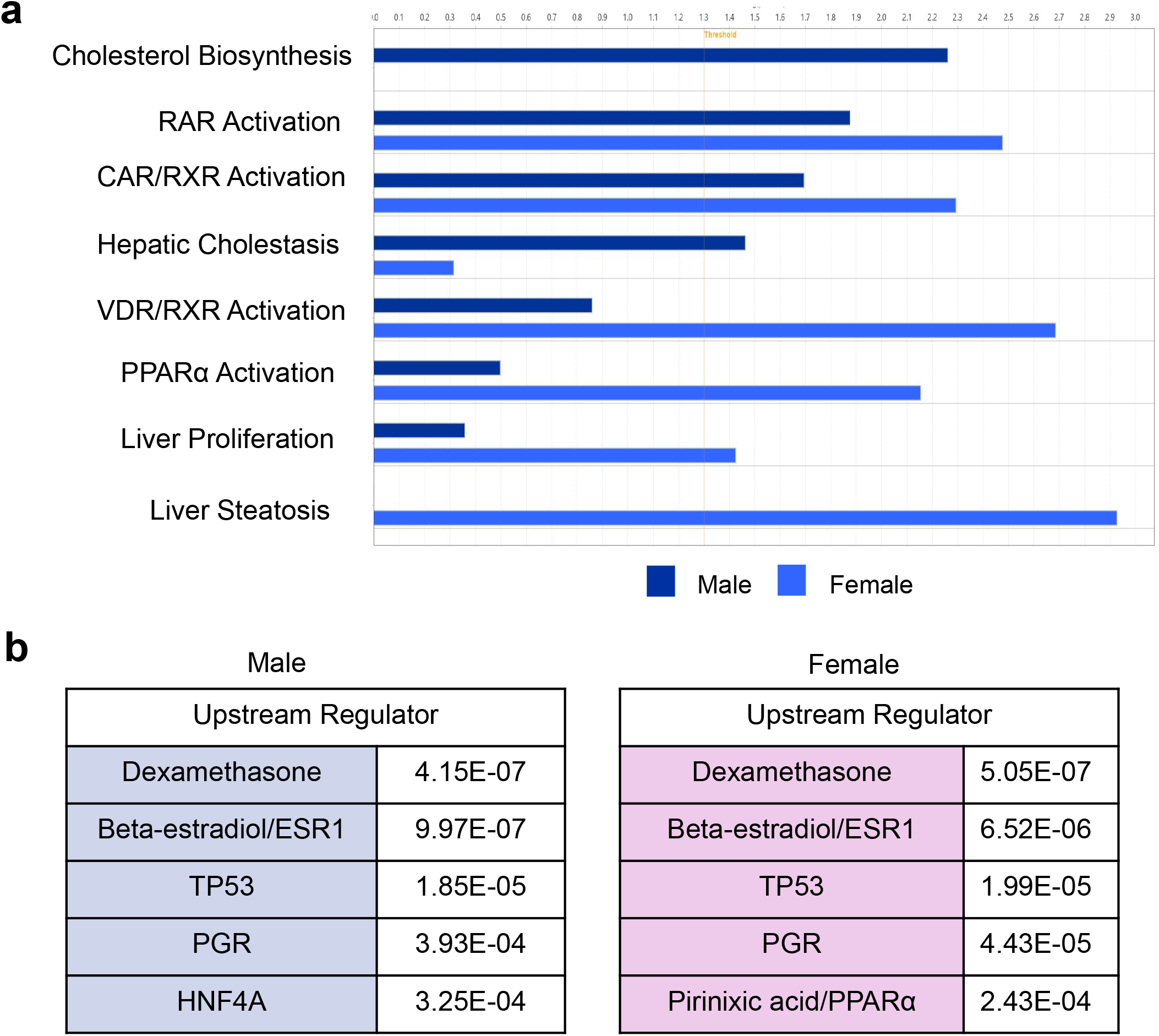
Nuclear receptors and p53 regulate FOXA2 targets in MASLD patients. **(a)** FOXA2 binding sites in male patients with severe steatosis and female patients with moderate steatosis were mapped to genes. IPA analysis identified activation of several nuclear receptors, including RAR, CAR in both males and females and VDR and PPARα only in females. (**b**) IPA analysis identified similar upstream regulators of genes bound by FOXA2 in female and male MASLD patients, including nuclear receptors ESR1, PGR, and dexamethasone/GR, as well as p53. In addition, PPARα regulates FOXA2 targets in livers of female MASLD patients, consistent with pathway analysis in (**a**).

We performed IPA upstream analysis to identify additional regulators of gernes bound by FOXA2. Interestingly, the regulators were similar between males and females, including nuclear receptors ESR1 and PGR, suggesting the gene expression in livers of male patients is feminized (**Fig. 2b**). In addition, PPARα regulates FOXA2 targets in livers of female MASLD patients, consistent with pathway analysis. Furthermore, upstream analysis identified p53 as a regulator of genes bound by FOXA2 in these individuals. Activation of p53 signaling we observe in livers of MASLD patients has also been reported in a laminopathy model, in livers of mice deficient in Zmpste24 protease which processes prelamin to mature lamin A. *Zmpste24* mutant mice have irregular nuclei and develop hepatic steatosis ^12^.

In addition, p53 is identified as a regulator of differentially expressed genes in MASLD patients (**Fig. 3c**). These genes, important to senescence and ageing, regulate cell cycle progression and include *Cdkn1a, Tbx3, Serpine1*, and *P4ha1. Cdkn1a* controls the G_1_ transition while *Tbx3* manages the G_2_-M transition. Activation of Cdhn1a and repression of Tbx3 as seen in our data (**Fig. 3c**), have been shown to result in an ageing and senescence phenotype when activated by p53^16,17^. Serpine1, upregulated in livers of MASLD patients, is also activated by p53, resulting in cell cycle arrest, DNA damage signaling, and increased pro-fibrosis markers^18^. Furthermore, suppression and silencing of Ph4a1 by p53, as observed in our RNA-Seq data, has been shown to decrease cell proliferation and cell cycle arrest in the G_1_ phase^19^. Activation of p53, as seen in MASLD patients (**Fig. 3c**) and *Zmpste24* mutant mice tie ZMPSTE24 and lamin A to our previously described model relating changes at the nuclear lamina to redistribution of FOXA2 binding and development of steatosis ^7^.

**Figure 3.**
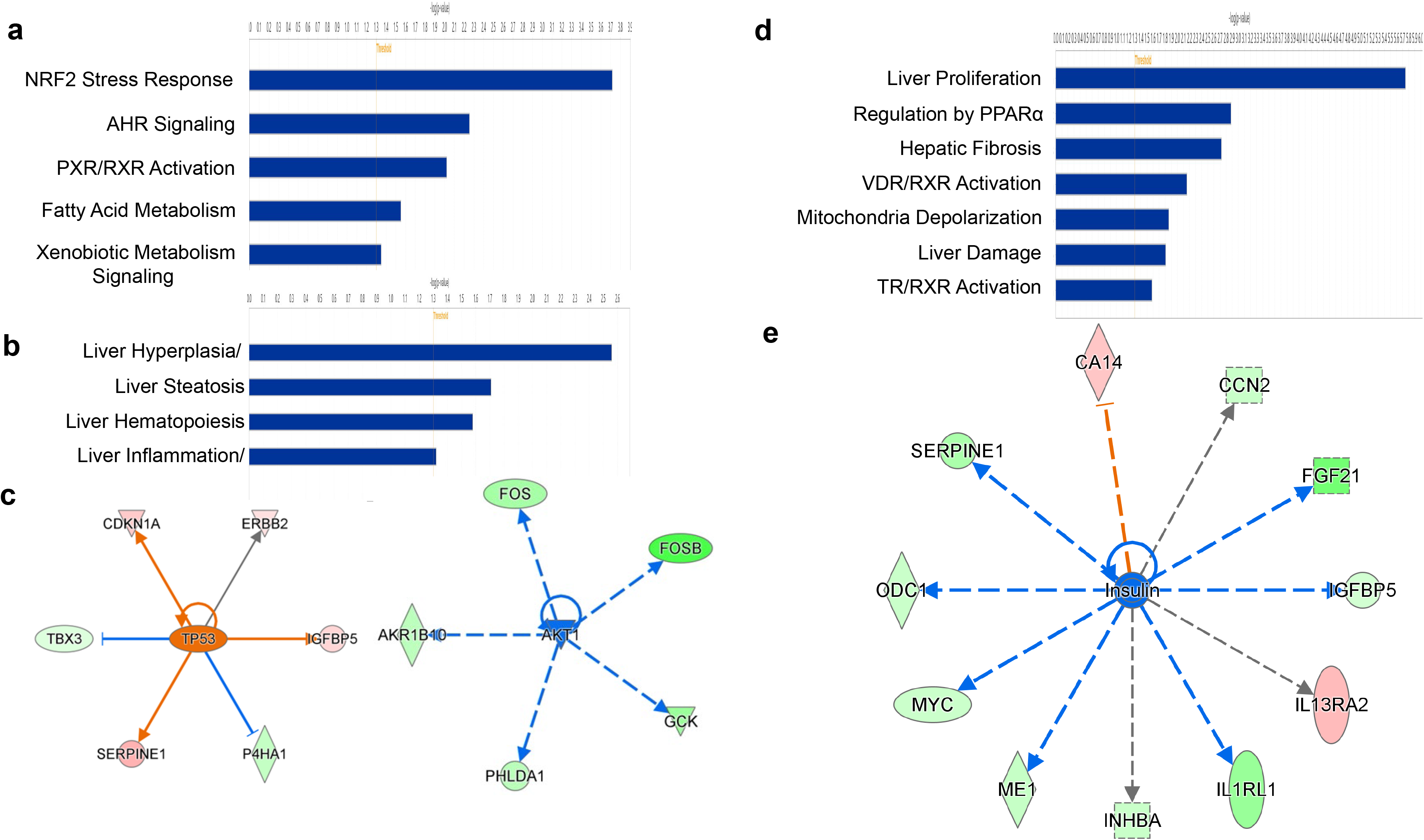
Differential gene expression analysis in male MASLD patients. (a) IPA analysis of differentially expressed genes in male MASLD patients (all with both moderate and severe steatosis identified pathways such as “NRF2 stress response”, “Ahr signaling” and activation of nuclear receptor PXR, (**b**) functions including liver steatosis and inflammation, (**c**) and networks demonstrating upregulated p53 activity (orange) and downregulated Akt1/insulin signaling (blue). (**d**) IPA analysis of differentially expressed genes in male MASLD patients with severe steatosis identified activation of several nuclear receptors (TR, VDR) and regulation by PPARα, as well as liver proliferation and damage pathways, (**e)** and a more extensive network of genes regulated by insulin. Activity of this regulator is inhibited (blue).

### Gene expression regulated by glucose and insulin is downregulated in MASLD patients

Typically, large cohorts of patients are needed to determine differential gene expression in a disease^20^. The livers we obtained from MASLD patients were well phenotyped depending on the level of triglyceride accumulation (mild, moderate, severe). Surprisingly, we were ablet to obtain a large number of differentially expressed from RNA-seq of MASLD livers (male: 3 normal vs. 6 steatotic, 74 genes; 3 normal vs 3 severe, 163 genes; female: 3 normal vs 6 steatotic, 59 genes; 3 normal vs 3 moderate, 266 genes, all FDR,<10). IPA analysis of differentially expressed genes in male MASLD patients (all with both moderate and severe steatosis) identified pathways such as “NRF2 stress response”, “Ahr signaling” and activation of nuclear receptor PXR, and liver functions including liver steatosis and inflammation (**Fig. 3a, b**). Networks corresponding to this gene expression included upregulated p53 activity (orange) and downregulated Akt1/insulin signaling (blue). IPA analysis of differentially expressed genes in male MASLD patients with severe steatosis identified activation of several nuclear receptors (TR, VDR) and regulation by PPARα, as well as liver proliferation and damage pathways, and a more extensive network of genes regulated by insulin (**Fig. 3d, e**) . Activity of this regulator is inhibited (blue).

Next, we performed IPA analysis of differentially expressed genes in female MASLD patients with moderate steatosis and identified activation of several nuclear receptors, including FXR, CAR, LXR, and PXR, as well cholesterol biosynthesis and fatty acid metabolism pathways (**Fig. 4a**). IPA network analysis of these genes demonstrated that activity of regulators, glucose and insulin, is inhibited (blue), while that of LXR (Nr1h3) is activated (orange) (**Fig. 4b**), consistent with cholesterol biosynthesis pathway identified in **Fig. 4a**. We observe downregulation of insulin signaling in both male and female MASLD patients with steatosis who have yet to progress to MASH. Insulin resistance is a hallmark of type II diabetes (T2D). Our genomic data suggest that although steatosis is considered benign, these patients are suffering from severe metabolic dysfunction. Comparison of overrepresented pathways in male MASLD patients with severe steatosis and female patients with moderate steatosis show showed sexual dimorphism. Although several pathways, such as liver proliferation, PPARα regulation and VDR activation are common, many, such as cholesterol biosynthesis and activation of nuclear receptors (FXR, CAR, LXR, PXR) is only observed in females.

**Figure 4.**
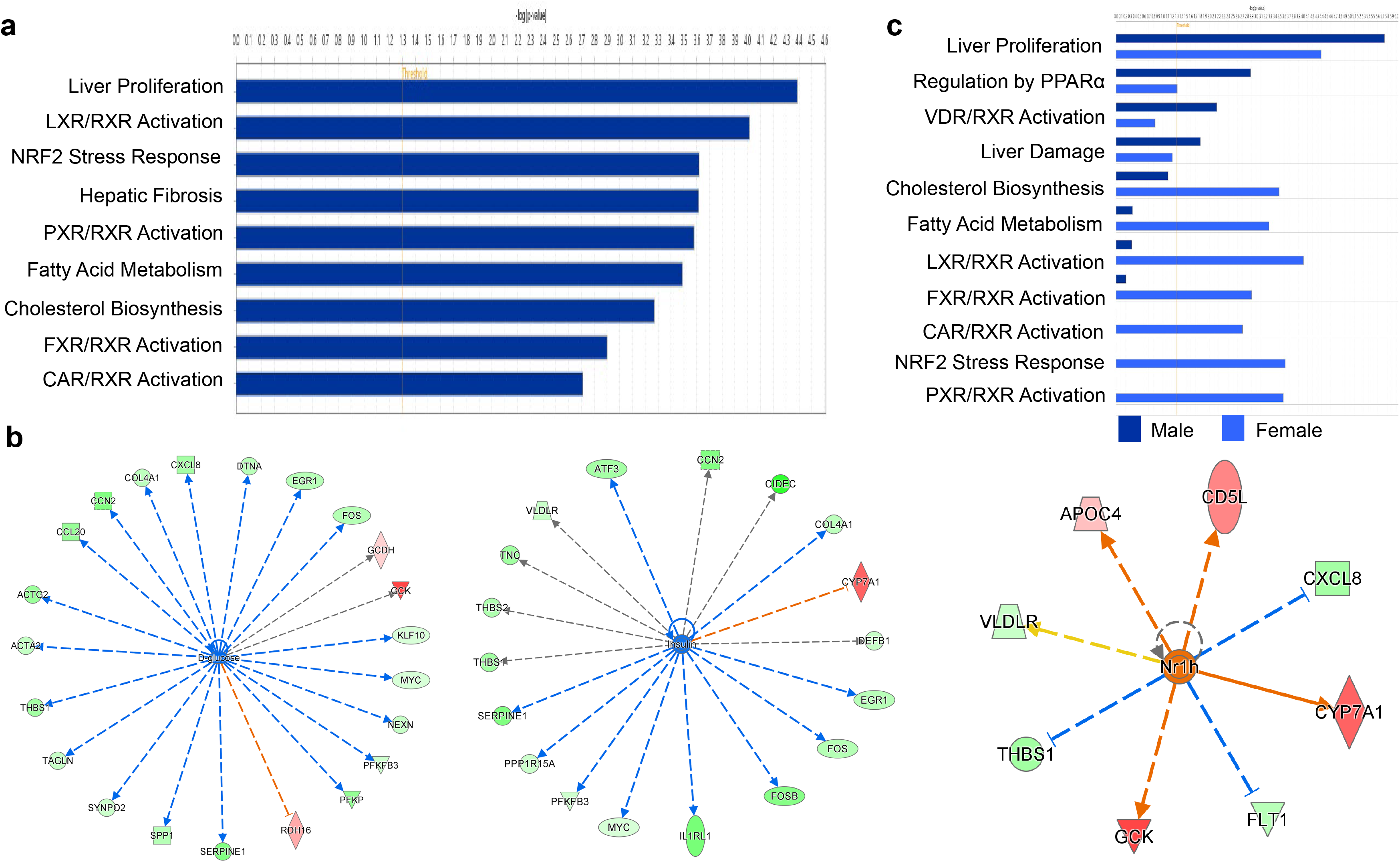
Differential gene expression analysis in female MASLD patients. (**a**) IPA analysis of differentially expressed genes in female MASLD patients with moderate steatosis identified activation of several nuclear receptors, including FXR, CAR, LXR, and PXR, as well cholesterol biosynthesis and fatty acid metabolism pathways. (**b**) IPA network analysis shows activity of regulators, glucose and insulin, is inhibited (blue), while that of LXR (Nr1h3) is activated (orange), consistent with cholesterol biosynthesis pathway identified in (**a**). (**c**) Comparison of overrepresented pathways in male MASLD patients with severe steatosis and female patients with moderate steatosis shows sexual dimorphism. Although several pathways, such as liver proliferation, PPARα regulation and VDR activation are common, many, such as cholesterol biosynthesis and activation of nuclear receptors (FXR, CAR, LXR, PXR) is only observed in females.

### Zmpste24 connects changes at the nuclear lamina to activated p53 signaling in MASLD patients

Activation of p53 signaling was consistent between Foxa2 targets and differentially expressed genes in livers of MASLD patients (**Fig. 2b, 3c**). Similarly, p53 activity is upregulated in livers of *Zmpstd24* mutant mice^12^. As previously stated, Zmpste24 processes prelamin into mature lamin A that can be used to maintain the proper shape of the nucleus and bind LADs^21^. Loss of proper Zmpste24 function results in improper nuclear shape and dysregulation of lamin binding, leading to improper Foxa2-dependent gene regulation and the development of steatosis ^11,12^. Hence, we proceeded to determine expression of ZMPSTE24 and prelamin in livers of MASLD patients. Indeed, there was a substantial prelamin accumulation in male and female patients with various degrees of steatosis (male: moderate, severe; female: mild, moderate) (**Fig.5**, top panel). These changes are caused by downregulation of ZMPSTE24 expression in livers of MASLD patients ((**Fig.5**, bottom panel)

**Figure 5.**
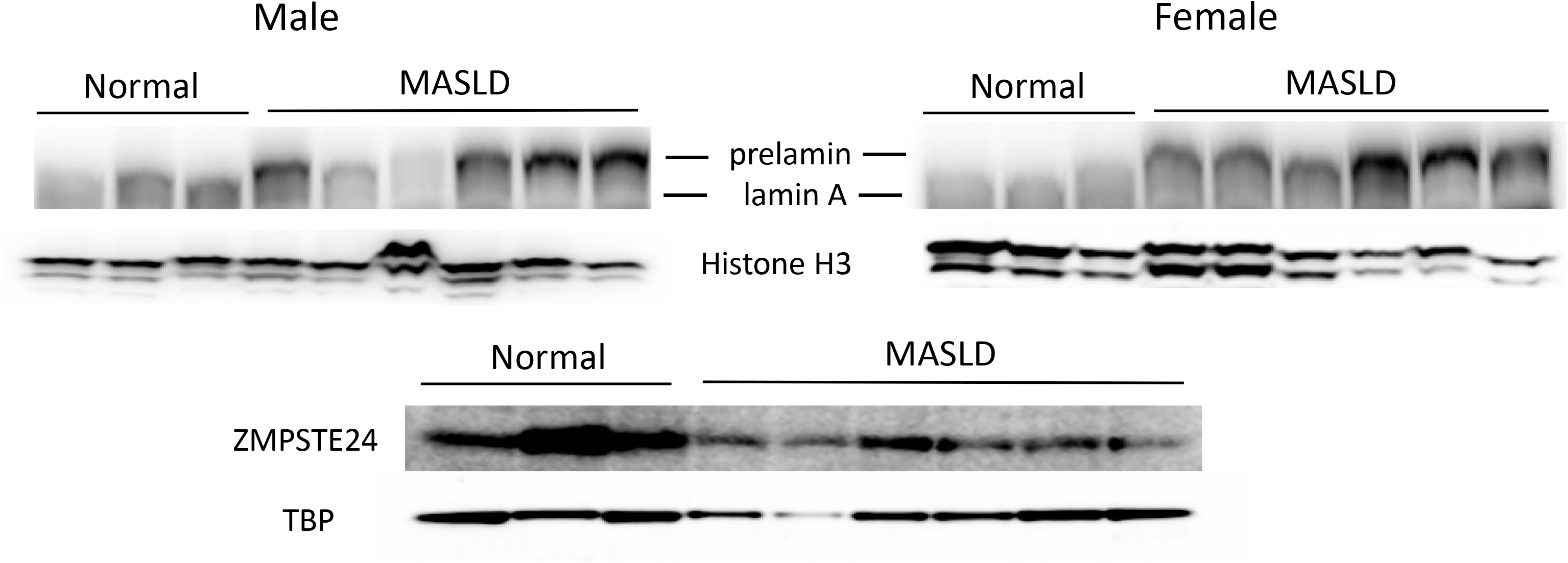
ZMPSTE24 downregulation and prelamin accumulation in MASLD patients. Western blot analysis of protein nuclear extracts from livers of three normal and six MASLD patients with antibodies to Lamin A and histone H3 (loading control, males top left panel, females top right panel. Prelamin accumulates in all MASLD patients. Same analysis with antibody to ZMPSTE24 and TBP( (loading control, bottom panel). ZMPSTE24 is downregulated in MASLD patients

Our results demonstrate that loss of Zmpste24 function is directly responsible for the development and progression of steatosis as prelamin accumulation leads to altered lamina shape (**Fig. 1a**), redistribution of LADs and loss of Lamin B1 signal (**Fig. 1b**) and loss of proper gene expression (**Fig. 3, 4**). This places Zmpste24 as a regulatory factor at the nuclear lamina that is upstream of our previously described Foxa2-dependent mechanism leading to severe steatosis. ^7^

This report extends our previous studies on a Foxa2-dependent mechanism leading to severe steatosis in aging & laminopathy mouse models and MASLD patients (**Fig. 6**). Loss of Zmpste24 leads to loss of prelamin processing and prelamin accumulation resulting in nuclear blebbing. Subsequently, LAD binding is disrupted and redistributed, leading to altered Foxa2 biding and development of steatosis ^7^.

**Figure 6.**
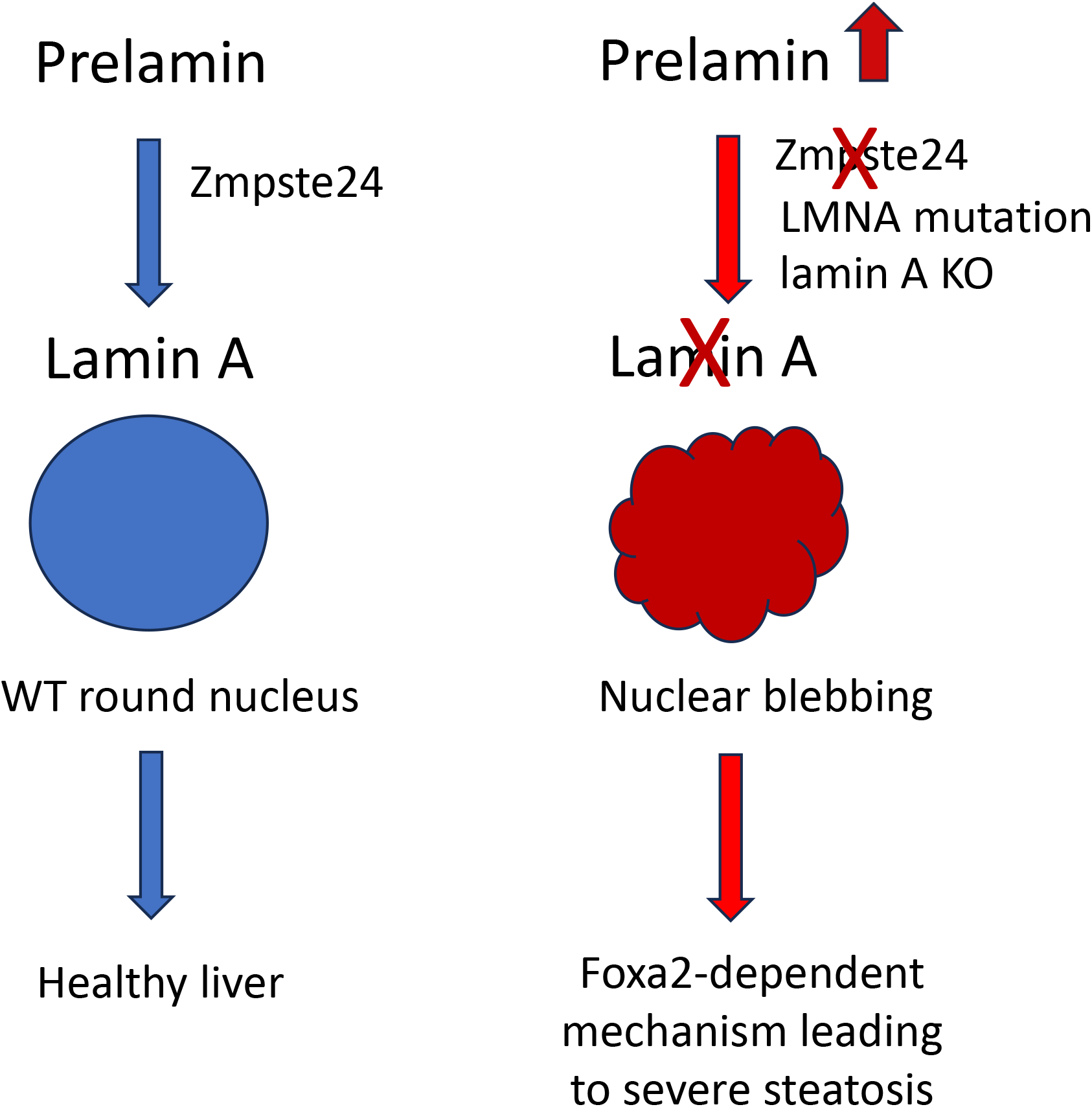
New mechanism upstream of Foxa2-dependent mechanism^7^ for development of MASLD. In healthy cells, prelamin is processed to mature lamin A by Zmpste24 enzyme. Normal lamin A expression is crucial to formation of proper round nuclei (left panel). Inhibition of Zmpste24 or LMNA mutation leads to accumulation of prelamin and no mature lamin A, leading to nuclear blebbing and dysmorphic nuclei found in MASLD patients^7^. Our results place Zmpste24 as a regulatory factor at the nuclear lamina that is upstream of our previously described Foxa2-dependent mechanism leading to severe steatosis.^7^.

## Discussion

No specific cause of MASLD has been established and currently there is no approved treatment for the disease. We and others have shown that nuclear shape changes in MASLD patients ^6,7^ and described a novel Foxa2-dependent mechanism for development of MASLD. Here, we extend our findings and demonstrate downregulation of ZMPSTE24 and accumulation of prelamin n both male and female MASLD patients, leading to the observed change in nuclear morphology. Hence, Zmpste24 is an upstream regulator that can alter LAD distribution and Foxa2 binding, dysregulation of which leads to development of steatosis in MASLD. A recent study of US persons with HIV on antiretroviral (AR) therapy found that nearly half have fatty liver, 90% of which is due to MASLD. A fifth of these individuals progress to liver fibrosis ^22^. Interestingly, antiretrovirals inhibit activity of ZMPSTE24, leading to accumulation of prelamin in cells of these patients ^23^. Hence, we propose that despite different etiology, the same fundamental mechanism involving inhibition of ZMPSTE24 and accumulation of prelamin that leads to abnormal nuclear morphology is involved in development of MASLD.

Zmpste24 cleaves farnesylated prelamin leading to mature lamin A. When Zmpste24 is inhibited, prelamin remains farnesylated. Farnesyltransferase inhibitor (FTI) Lonafarnib has been approved for treatment of Hutchinson-Giford progeria syndrome, caused by accumulation of farnesylated prelamin, and other laminopathies. Use of Lonafarnib and another FTI Tipifarnib has been shown to improve metabolic syndrome induced by AR therapy in mice ^24^. We suggest using FTI inhibitors for treating MASLD in patients. These medications should reduce prelamin accumulation and re-establish proper nuclear shape, eliminating downstream effects leading to development of steatosis and further progression to Metabolic dysfunction-associated steatohepatitis MASH.

We also find that genes regulated by glucose and insulin are downregulated in MASLD patients. Reduced insulin sensitivity and dysregulated glucose metabolism are hallmarks of T2D. Proper glycemic control can be maintained by the liver through glucose production (gluconeogenesis) and glycogen breakdown (glycogenolysis) under normal conditions. The liver also helps maintain proper lipid metabolism and fatty acid storage and release, and the disruption of these functions also leads to insulin resistance. Thus, it is important to recognize severe metabolic dysfunction in MASLD patients and need for treatment before the disease progresses to MASH.

## Methods

### Human studies

Liver tissue from non-steatotic control (hepatic TG< 5%) and MASLD donors were obtained through Sekisui Xenotech Tissue Biobank. Criteria for samples: 1) individuals younger than 60 since lamina shape changes occur in old individuals, 2) assurance that individuals were not heavy drinkers. Based on the criteria, we were able to obtain 3 samples per each category. In total, 9 samples from female patients (3 control, 3 mild MASLD, 3 moderate MASLD) and 9 samples from male patients (3 control, 3 mild/moderate MASLD, 3 severe MASLD) were procured. Female donors were 17–59yr old (healthy, 17–59; MASLD, 31–47) Male donors were 21–51yr old (healthy, 26–45; MASLD, 21–51).

### Chromatin immunoprecipitation and ChIP-Seq

Snap-frozen mouse or human liver (100 mg) was used to prepare chromatin. ChIP and ChIP-seq were performed as reported previously^25^. Briefly, liver tissue was minced in cold PBS and cross-linked in 1% formaldehyde/PBS for 15 min with constant rotation in a Labquake tube rotator. Cross-linking was quenched by adding glycine to a final concentration of 0.125M. Nuclear lysate was sonicated using a Diagenode Bioruptor Pico for 14 cycles (30 sec on/30 sec off). FOXA2-specific rabbit antiserum (Seven Hills Bioreagents WRAB-1200) and rabbit antibody to lamin B1 (Abcam Ab16048), were used for immunoprecipitation. Libraries were made according to standard Illumina protocol (end-repair of ChIP DNA, addition of A base to the 3′-ends, adapter ligation, and amplification). We used multiplex adapters for sequencing and Kapa HiFi DNA polymerase (Kapa Biosystems) for PCR amplification (16 cycles). Library fragments were isolated using Pippin HT agarose gel. The purified DNA was captured on an Illumina flow cell for cluster generation. Libraries were sequenced on Illumina NextSeq 500 and NextSeq 2000 instruments following the manufacturer’s protocols.

### Protein analysis

Protein extracts preparation and protein immunoblot analysis were performed as reported previously^13^. The primary antibodies used were rabbit antibody to histone H3 (Cell Signaling 4499, 1:4000), mouse antibody to LAMIN A/C (Cell Signaling mAb 4777, 1:2000), rabbit antibody to ZMPSTE24 (Invitrogen PA1-16965, 1:2000), and rabbit antibody to TBP (Abcam ab220788, 1:2500).

### RNA isolation, analysis, and sequencing

Liver RNA was isolated from heterozygous liver donor tissue as described previously^25^. The quality of the RNA samples was analyzed using an Agilent RNA 6000 nano kit (Bioanalyzer, Agilent Technologies). Samples with RIN scores above 9.5 were used in library preparation. One microgram of total RNA was used to isolate mRNA (NEBNext Poly(A) mRNA magnetic isolation module). Libraries of the resulting mRNA were prepared using a NEBNext Ultra II RNA library preparation kit. All samples were sequenced on an Illumina NextSeq 2000. Three replicates were sequenced for each category (female: Normal, Mild, Moderate; male: Normal, Moderate, Severe).

### ChIP-Seq analysis

Reads were aligned to human genome (hg19; NCBI build 37) using BWA v0.7.12 (Li and Durbin 2009). Duplicate reads were removed using Picard v 1.134 (http://picard.sourceforge.net). Reads (Phred score ≥ 30) that aligned uniquely were used for subsequent analysis. Data from three human patients were merged for FOXA2 and lamin B1 binding (three normal and three MASLD). Epic peak caller^26^ (https://github.com/biocore-ntnu/epic2) was used to determine lamin B1–associated domains in human patients (hg19 as species, window size of 10 kb, gap size of three, FDR 5%). PeakSeq^27^ was used to identify FOXA2 bound peaks against input controls (FDR 5%, Q-value < 0.07).

### RNA-Seq analysi**s**

RNA-Seq reads were aligned using STAR^28^ to human genome (hg19; NCBI build 37). Expression levels were calculated using RSEM^29^. Differential expression analysis of RNA-seq (P-value < 0.05) was performed in R using edgeR package)^15^ with a Benjamini–Hochberg FDR of 5%.

### Functional analysis

Heatmaps of ChIP-seq coverage were generated by deepTools^30^. ChIP-seq peaks were associated with closest genes using GREAT (McLean et al. 2010), which were subsequently used for Ingenuity Pathway Analysis (pathway and upstream analysis). IPA was also used for pathway, network, and upstream analysis of differentially expressed genes.

## Data Availability

All genomic sequencing data generated in this study have been submitted to the NCBI Gene Expression Omnibus (GEO; https://www.ncbi.nlm.nih.gov/geo/) under accession number GSEXXX

## Acknowledgments

We thank M. Murphy, N. Reddy, and X. Wei for technical assistance

